# Transcription factors regulating the fate and developmental potential of a multipotent progenitor in *C. elegans*

**DOI:** 10.1101/2022.07.01.498512

**Authors:** Evan M. Soukup, Jill C. Bettinger, Laura D. Mathies

**Author notes:** To whom correspondence should be addressed: Laura Mathies Department of Pharmacology and Toxicology Box 980613 Richmond, VA 23298-0613.

## Abstract

Multipotent stem and progenitor cells have the capacity to generate a limited array of related cell types. The two *C. elegans* somatic gonadal precursors (SGPs) are multipotent progenitors that generate all 143 cells of the somatic gonad, including complex tissues and specialized signaling cells. To screen for candidate regulators of cell fate and multipotency, we identified transcription factor genes with higher expression in SGPs than in their differentiated sister, the head mesodermal cell. We used RNA interference or genetic mutants to reduce the function of 183 of these genes and examined the worms at two developmental stages. Eight genes were identified that regulate the SGP fate, including the *Ci/GLI* homolog *tra-1,* which is the terminal regulator of sex determination. We show that *tra-1* regulates SGP fate independently of its role in sex determination, supporting the idea that *tra-1* retains ancestral functions. Four genes were necessary for SGPs to generate the correct number and type of descendant cells. We show that the *E2F* homolog, *efl-3,* regulates the cell fate decision between distal tip cells and the sheath/spermathecal precursor. We find that the FACT complex gene *hmg-4* is required for the generation of the correct number of SGP descendants, and we define an earlier role for the *nhr-25* nuclear hormone receptor-encoding gene, in addition to its previously described role in regulating the asymmetric division of SGPs. Overall, our data show that genes regulating cell fate are largely different from genes regulating developmental potential, demonstrating that these processes are genetically separable.

## Introduction

As animals develop from a single-celled zygote, their cells transition through different states of developmental potential. Pluripotency is the capacity to generate all cell types of the mature organism. The expression of cocktails of transcription factors, called pluripotency factors, can induce differentiated cells to become pluripotent. The resulting induced pluripotent stem cells (iPSCs) hold great therapeutic potential because they can be patient-derived and have few ethical concerns (Deinsberger et al., 2020).

Adult stem cells and lineage restricted progenitors are multipotent, meaning that they can generate a more limited array of cell types derived from a single lineage.

Considerably less is known about the regulation of multipotency. One of the best-characterized multipotent stem cells is the hematopoietic stem cell (HSC), which gives rise to all blood cell types. Although differentiated blood cells have been converted into multipotent HSCs (Riddell et al., 2014), efforts to induce HSCs from unrelated cell types have been less successful (Esposito, 2016). Genetic studies in model systems may reveal conserved strategies that give multipotent progenitors the capacity to produce diverse but restricted cell types.

The *C. elegans* somatic gonadal precursors (SGPs) are multipotent progenitors that generate all somatic cells of the gonad. In hermaphrodites, two SGPs produce 143 cells through two periods of cell division (Kimble and Hirsh, 1979). During the first and second larval stages, each SGP generates six daughter cells – one distal tip cell (DTC), two sheath/spermathecal precursors (SS), one dorsal uterine precursor (DU), one ventral uterine precursor (VU), and one cell with the potential to become the anchor cell (AC). One of two bipotential AC/VU cells will differentiate as the AC, the other will become a VU precursor; this decision is determined by lateral signaling between the two AC/VU cells (Greenwald et al., 1983; Kimble and Hirsh, 1979; Seydoux and Greenwald, 1989). The DTCs and AC differentiate into specialized signaling cells: the AC induces formation of the vulva (Kimble, 1981) and two DTCs signal to the underlying germ cells to promote their proliferation (Kimble and Hirsh, 1979; Kimble and White, 1981). The remaining progenitors divide again during the third and fourth larval stages to produce 140 cells. These cells form diverse tissues such as a contractile sheath that is important for ovulation, and the spermatheca and uterus, which are tubular epithelia that house sperm and developing embryos, respectively (reviewed in Hubbard and Greenstein, 2000). Thus, the *C. elegans* SGPs produce diverse cell and tissue types through a completely defined cell lineage.

We have used the SGP/hmc cell fate decision as a paradigm for understanding how multipotency is regulated during development. The sisters of the SGPs are the left and right head mesodermal cells (hmcL and hmcR). These cells do not divide further; instead hmcR undergoes programmed cell death and hmcL terminally differentiates as the single head mesodermal cell (hmc), a cell of unknown function and neuron-like morphology (Altun and Hall, 2009; Sulston et al., 1983). Therefore, the cell fate decision between SGPs and hmcs is one between multipotency and differentiation. We analyzed the transcriptomes of SGPs and hmcs and identified genes that are differentially expressed between these two cell types (Mathies et al., 2019). Of particular interest to this study are genes with higher expression in SGPs than hmcs, referred to here as SGP-biased genes; among these were 184 genes encoding transcription factors (TFs). One might expect that SGP-biased TFs will include regulators of SGP fate and multipotency.

We previously identified four genes that are important for the SGP cell fate.

*hnd-1* encodes a bHLH transcription factor orthologous to *dHand* in vertebrates (Mathies et al., 2003), and three genes, *pbrm-1, swsn-1,* and *swsn-4,* encode subunits of a SWI/SNF chromatin remodeling complex (Large and Mathies, 2014). *hnd-1* or SWI/SNF mutants have SGPs that express markers of the SGP and hmc cell fates, and they are frequently missing one or both gonadal arms in the adult (Large and Mathies, 2014; Mathies *et al*., 2003). These mutant phenotypes suggest that SGPs are partially transformed into hmcs, and, as a result, they fail to produce mature gonadal cell types. Importantly, null mutations resulted in incompletely penetrant phenotypes, strongly arguing for redundancy in the regulation of the SGP/hmc cell fate decision.

To comprehensively search for regulators of SGP fate and multipotency, we examined the function of 183 SGP-biased TF genes using RNAi. Eight genes are important for normal expression of cell fate markers in SGPs, indicating that they play a role in the SGP/hmc cell fate decision. One such gene is the sex determination gene *tra-1* (Hodgkin, 1987; Zarkower and Hodgkin, 1993); we show that *tra-1*’s function in the SGP/hmc cell fate decision is independent of its sex determining function. Four genes are necessary for the production of the correct number and type of SGP descendants at the L3 larval stage; these genes are outstanding candidates for the regulators of multipotency in SGPs. We describe an earlier function for the nuclear hormone receptor gene *nhr-25,* in addition to its role in regulating the asymmetric division of SGPs (Asahina et al., 2006), and we identify new roles for the FACT complex gene *hmg-4* and the E2F gene *efl-3* in somatic gonad development. Importantly, the genes regulating cell fate and multipotency are largely non-overlapping, indicating that the genetic controls of cell fate and multipotency are distinct.

## Materials and Methods

### Strains

Animals were grown at 20°C. All strains were derivatives of Bristol strain N2 (Sulston and Horvitz, 1977). The following alleles and transgenes were used in this study and are described in *C. elegans II* (Hodgkin, 1997) or cited references. *LGI: rdIs7* [*hnd-1::rde-1*] (Large and Mathies, 2014), *arTi361* [*gfp(flexon)::h2b*] (Shaffer and Greenwald, 2022). *LGII: rdIs35* [*ehn-3::tdTomato*] (Mathies *et al*., 2019), *ccIs4444* [*arg- 1::GFP*] (Kostas and Fire, 2002), *znfx-1(gg561)* (Wan et al., 2018). *LGIII*: *tra-1(e1099)*, *unc-119(ed3)* (Maduro and Pilgrim, 1995). *LGIV*: *lin-22(ot287)* (Doitsidou and Hobert, 2019). *LGV*: *rde-1(ne219)* (Tabara et al., 1999), *ceh-75(gk681)* (Consortium, 2012), *him-5(e1490), qIs70* [*lag-2::YFP*] (Kidd et al., 2005), *syIs51 [cdh-3::CFP]* (Inoue et al., 2002). *LGX*: *arTi237* [*ckb-3p::Cre*] (Shaffer and Greenwald, 2022). Dominant GFP balancer for *LGI* and *LGIII*: *hT2[qIs48]*.

### SGP-biased transcription factors

We queried the list of 2749 SGP-biased genes (FDR ≤ 0.01, fold change ≥ 2) (Mathies *et al*., 2019) against the worm transcription factor databases (Reece-Hoyes et al., 2005; Reece-Hoyes et al., 2011) and included any gene that was present in either database. In total, 184 of the SGP-biased genes were predicted to encode transcription factors (File S1). Three of these genes*, lin-22, ceh-75,* and *znfx-1*, did not have available RNAi clones. We obtained loss-of-function alleles for each of these genes and crossed *rdIs35* [*ehn-3::tdTomato*] and *ccIs4444* [*arg-1::GFP*] into the mutant backgrounds. *znfx-1* is located on the same chromosome as *rdIs35* [*ehn^-^3::tdTomato*] and *ccIs4444* [*arg-1::GFP*]. We attempted to recombine the reporters with *znfx-1(gg561)* and were unsuccessful; therefore, this allele was only screened in our L4 screen. The only gene we were unable to screen was *nhr-192* because the RNAi clone did not grow and there were no loss-of-function alleles affecting only this gene. In total, we screened 183 of the 184 predicted SGP-biased transcription factors.

### Feeding RNAi

RNAi by feeding was performed essentially as described in Kamath and Ahringer (Kamath et al., 2001). RNAi clones were obtained from a nearly complete transcription factor RNAi library (MacNeil et al., 2015) or the commercially available RNAi libraries (Kamath et al., 2003; Rual et al., 2004). The clones were first streaked onto LB plates containing 50 μg/mL carbenicillin and 12.5 μg/mL tetracycline. Liquid LB cultures containing 50 μg/mL carbenicillin were inoculated from a single colony and grown for 16-18 hours at 37°C. Nematode growth medium (NGM) plates containing 25 μg/mL carbenicillin and 1 mM IPTG were seeded with 150 μL of bacterial culture and incubated at 20°C for 2-3 days before worms were placed on the plates. Any RNAi clone that produced a phenotype was sequenced to ascertain that the clone was correct. Two clones were found to be incorrect; they were replaced with clones from other RNAi libraries and screened again.

### L1 screen

L4 staged worms were placed on the RNAi plates and allowed to develop for 36-48 hours. The resulting adult worms were transferred to new RNAi plates and allowed to lay eggs for one hour. Two collection windows were generated for each RNAi. The F1 progeny were screened approximately 24 hours later, when they were mid-L1 staged larvae, using fluorescence and differential interference contrast (DIC) microscopy. Approximately 20 L1 staged worms were observed for the initial screen.

The empty RNAi vector control was included with every batch. We recorded the number of cells in the gonad, expression of tdTomato in SGPs or anywhere outside of the gonad, and expression of GFP fluorescence in the gonad and the hmc. For GFP, we noted any expression that was brighter than that observed in the empty vector control. We also noted any cellular morphology changes. We performed a secondary screen for any genes having: 1) ≥ 25% of SGPs with GFP expression, 2) GFP expression in SGPs that was brighter than the control, or 3) any difference in the number of tdTomato-expressing cells in the gonad. We followed a similar protocol for the secondary screen, except we scored a minimum number of 40 L1 staged worms, and we noted three levels of GFP expression in the SGPs – dim, distinct, or bright.

### L4 screen

L4 staged worms were placed on the RNAi plates and allowed to develop for 36-48 hours. The resulting adult worms were transferred to new RNAi plates and allowed to lay eggs for ∼24 hours. The F1 progeny were screened as L4 larvae using a dissecting microscope. At least 50 worms were examined for gonadogenesis defects. We scored missing gonadal arms (one-arm, OA), disorganized gonads with a central patch of gonadal tissue (white patch, WP), and absent gonads (gonadless, Gon). Non-gonadal phenotypes were noted if observed. We performed a secondary screen using GS9401: *arTi361* [*gfp(flexon)::h2b*]; *arTi237* [*ckb-3p::Cre*] to mark all SGP descendants (Shaffer and Greenwald, 2022). RNAi was performed as for the L4 screen, except the adult worms were allowed to lay eggs for one hour and two collection windows were taken. The F1 progeny were screened approximately 36 hours later, at or shortly after the L2/L3 molt, using fluorescence and DIC microscopy. Approximately 50 worms were scored for each RNAi knockdown and the number and organization of SGP descendants was recorded. We performed a follow up screen on *efl-3* and *nhr-25* RNAi using markers for DTCs and the AC. For this purpose, we recombined *qIs70[lag-2::YFP]* and *syIs51[cdh-3::CFP]* to make RA344; this strain expresses YFP in DTCs and CFP in the AC. RNAi was performed as for GS9401, except the worms were allowed to develop for 48 hours and were examined as early L4 larvae.

### Tissue-specific RNAi

For RNAi clones that produced an embryonic or larval lethal phenotype, we repeated the RNAi using a tissue-specific RNAi strain containing *rde-1(ne219)* and *rdIs7* [*hnd-1::rde-1*]. This strain rescues *rde-1* in mesodermal lineages including SGPs (Large and Mathies, 2014). We crossed *rdIs35* [*ehn-3::tdTomato*] and *ccIs4444* [*arg-1::GFP*] into the tissue-specific RNAi background to create RA701. We crossed *arTi361* [*gfp(flexon)::h2b*] and *arTi237* [*ckb-3p::Cre*] into the tissue-specific RNAi background to create RA708. RNAi was performed as described above.

### sex-1, tra-1 and XO males

To examine XO males, we used a strain, RA481, that contains *rdIs35* [*ehn-3::tdTomato*] and *ccIs4444* [*arg-1::GFP*] in the *him-5(e1490)* background. The *him-5* mutation results in a higher rate of X chromosome nondisjunction and produces a higher incidence of males (Him) phenotype (Hodgkin et al., 1979). Males were distinguished from hermaphrodites by the size of the B cell nucleus. We crossed *rdIs35* [*ehn-3::tdTomato*] and *ccIs4444* [*arg-1::GFP*] into *tra-1(e1490)* and *sex-1(y263)* to create RA644 or RA705, respectively. *tra-1(e1490)* was balanced by the GFP-marked balancer *hT2[qIs48]*. *tra-1(e1490)* homozygotes were identified by the absence of GFP expression in the pharynx.

### Microscopy

Fluorescent reporters were visualized using an Axio Imager A1 microscope with a 63x Plan Apochromatic objective (Zeiss). Images were captured using an AxioCam MRm monochromatic camera with Zen software (Zeiss).

### Statistical analysis

Graphs were generated and statistical analysis was performed using Prism 9 version 9.3.1 (Graphpad). The number of SGP descendants in each RNAi knockdown was compared to the control RNAi using one-way ANOVA with Dunnett’s post-hoc multiple comparisons test.

## Results

We used RNAi to examine the function of 183 (of 184) predicted transcription factor genes with higher expression in SGPs than hmcs (Mathies *et al*., 2019). We took a two-pronged approach to screening. First, to identify genes that influence the SGP/hmc cell fate decision, L1 larvae were screened for differences in expression of SGP and hmc cell fate markers using fluorescence and DIC microscopy. Second, to identify genes that regulate SGP multipotency, L4 stage larvae were screened for gonadogenesis defects using a dissecting microscope. We reasoned that RNAi knockdown of genes required for multipotency would result in fewer or different gonadal cell types, which might lead to gross changes in gonadal morphology. For any genes that produced a lethal phenotype, the RNAi knockdown was repeated using an engineered tissue-specific RNAi strain that restricts RNAi to mesodermal lineages (Large and Mathies, 2014). For genes without an RNAi clone, we examined available genetic mutants, where possible (see Materials and Methods).

### L1 RNAi screen identifies candidate SGP fate determinants

We used a dual-color marker strategy to monitor the SGP and hmc cell fates (Figure 1A). In wild-type worms at the L1 stage, *ehn-3::tdTomato* is expressed exclusively in the two SGPs (Mathies *et al*., 2019) and *arg-1::GFP* is expressed in hmcs and enteric muscles in the tail (Kostas and Fire, 2002; Zhao et al., 2007). The hmc has a distinctive H-shaped morphology and location (Altun and Hall, 2009), making it easy to assess expression of GFP in hmcs. Since *arg-1::GFP* is expressed at low levels in SGPs (Large and Mathies, 2014), we determined the best developmental window for scoring the SGP fate. We found that *arg-1::GFP* expression diminished over time such that worms with six cells in the gonad had minimal GFP expression in SGPs (Figure 1B-C).

**Figure 1.**
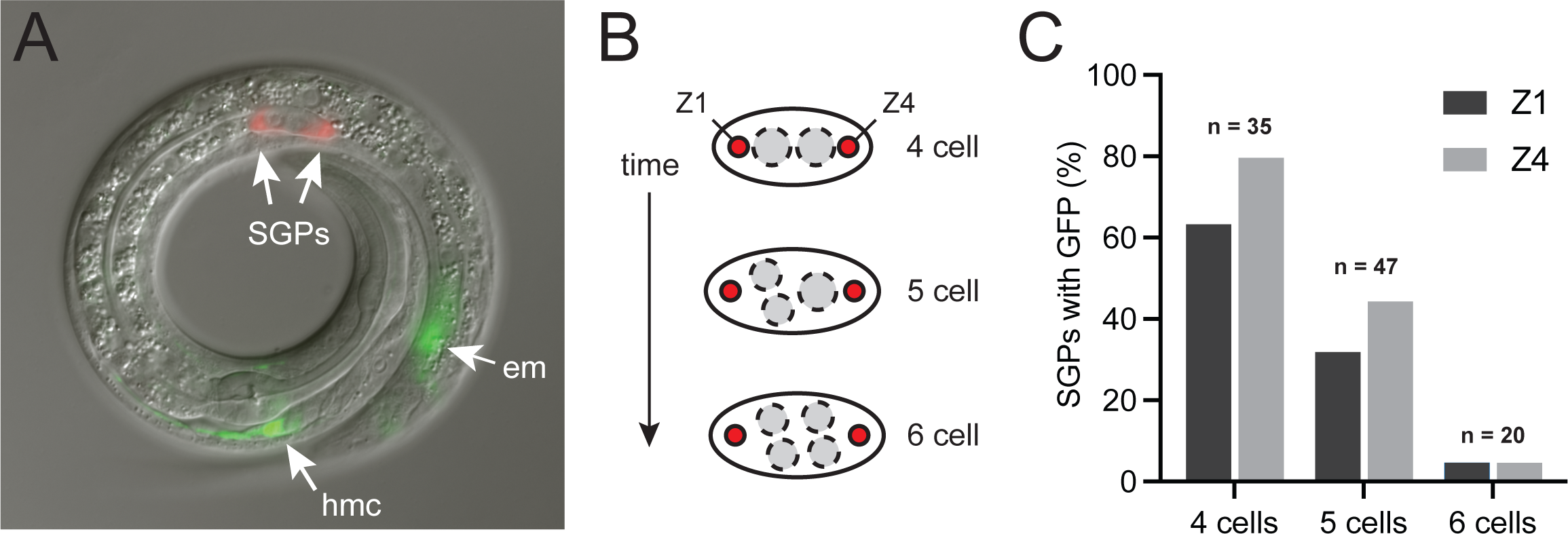
SGP and hmc markers. (A) Expression of *arg-1::GFP* (green) and *ehn-3::tdTomato* (red) in L1 animals. Fluorescent image overlay onto the corresponding DIC image. *ehn-3::tdTomato* is expressed exclusively in two somatic gonadal precursors (SGPs) within the gonad primordium; *arg-1::GFP* is expressed in the head mesodermal cell (hmc), an H-shaped cell near the pharynx in the head, and in enteric muscles (em) in the tail. (B) Diagram of the gonad primordium over time. L1 worms hatch with a 4-celled gonad primordium containing two SGPs (red) and two primordial germ cells (PGCs, gray). The PGCs divide before either SGP divides, resulting in four to six cells in the gonad. (C) Expression of *arg-1::GFP* in SGPs (Z1 and Z4) in wild-type worms with four to six cells in the gonad primordium.

We performed feeding RNAi on all of the SGP-biased transcription factors for which there were available RNAi clones (see Materials and Methods). In order to minimize the background level of *arg-1::GFP* expression, only worms with five or six cells in the gonad were scored; most had six cells. First, we assessed expression of *ehn-3::tdTomato.* Control animals almost always had two cells expressing tdTomato in the gonad primordium (*n*=582/583 worms), as did most of the RNAi knockdowns. We never observed *ehn-3::tdTomato* expression in hmcs or anywhere outside of the gonad primordium. There were 12 genes for which RNAi knockdown resulted in occasional worms with fewer than two tdTomato-expressing cells in the gonad (Table 1). These are candidate SGP fate determinants.

**Table 1.** SGP-biased TFs with altered SGP marker expression.

| gene | tdTomato expression in SGPs |  |
| --- | --- | --- |
|  | No expression (%) | SGPs (n) |
| *nhr-92* | 12% | 34 |
| *tra-1* | 6% | 34 |
| *R05D3.3* | 6% | 34 |
| *F44E2.7* | 6% | 34 |
| *icd-2* | 6% | 36 |
| *moe-3* | 5% | 44 |
| *fkh-6* | 3% | 30 |
| *nhr-136* | 3% | 34 |
| *K11D12.12* | 3% | 38 |
| *ceh-63* | 3% | 40 |
| *zip-9* | 2% | 42 |
| *ztf-23* | 2% | 42 |

Next, we examined expression of *arg-1::GFP*. We always observed GFP expression in a cell with the correct location and morphology to be the hmc. Control RNAi worms also had dim GFP expression in 4.1% of Z1 cells and 5.7% of Z4 cells (*n*=583 worms). Many of the RNAi knockdowns resulted in a higher percentage of SGPs expressing GFP. We chose 25% as a cutoff because it reflected two standard deviations from the mean for the control (File S1). Using this criterion, we identified 19 genes that, when inactivated, resulted in a higher percentage of SGPs expressing GFP or higher levels of GFP expression in at least one SGP (Table 2). These are candidate regulators of the SGP/hmc cell fate decision. In total, our primary screen identified 28 candidate genes.

**Table 2.**
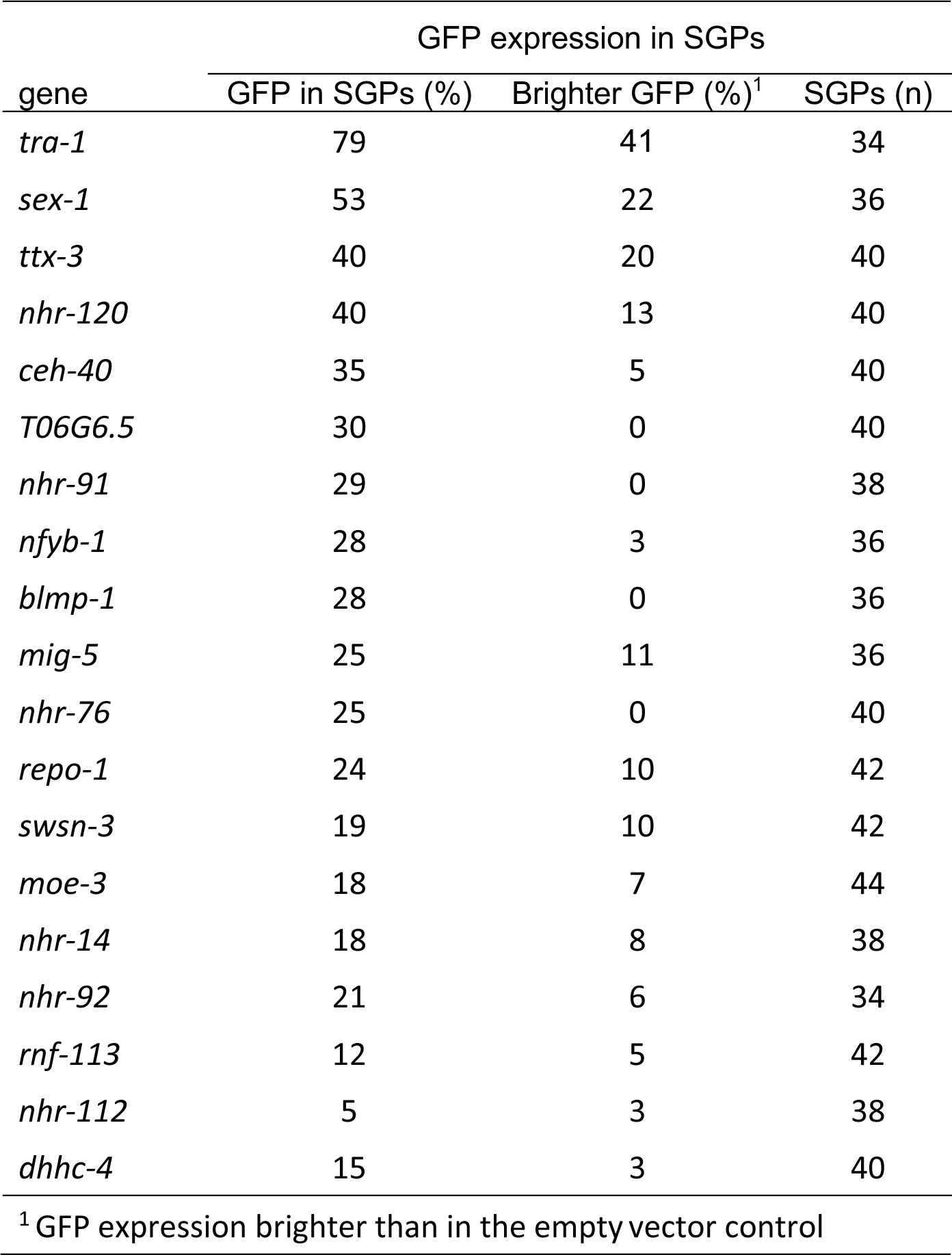
SGP-biased TFs with altered hmc marker expression.

Interestingly, two of our top candidate genes were members of the sex determination pathway. In *C. elegans,* sex is determined by the ratio of X chromosomes to autosomes, with XX individuals developing as hermaphrodites and XO individuals developing as males (Figure 2A). *tra-1* is the terminal regulator of somatic sex determination; it is required for hermaphrodite development (Hodgkin, 1987; Zarkower, 2006). *sex-1* is one of several X chromosome signal elements required to promote hermaphrodite development (Carmi et al., 1998). Loss-of-function mutations in *tra-1* or *sex-1* result in XX animals developing as males. *sex-1* also controls dosage compensation; therefore the transformed XX animals frequently die because of inappropriate X chromosome gene regulation (Carmi *et al*., 1998; Gladden et al., 2007). We examined genetic mutants of *tra-1* and *sex-1* using the SGP and hmc markers and found that the strong loss-of-function mutation *tra-1(e1490)* caused significant hmc marker expression in SGPs, in agreement with our RNAi result, while the viable *sex-1(y263)* allele did not (Figure 2B). *sex-1(y263)* is a splice acceptor mutation that affects sex determination and dosage compensation; it is not a null allele (Gladden *et al*., 2007). Thus, it is possible that RNAi produces a stronger phenotype and reveals a role for *sex-1* that is not seen in *sex-1(y263)*. We hypothesized that *tra-1* and *sex-1* loss of function might result in increased *arg-1::GFP* expression in SGPs because of the transformation of the animals toward the male fate. If this were true, we would expect XO males to express *arg-1::GFP* in SGPs. To test this idea, we examined *arg-1::GFP* expression in XO males and found that it is qualitatively similar to that of XX hermaphrodites (Figure 2B), suggesting that *tra-1* and possibly *sex-1* regulate the SGP cell fate decision independent of their role in sex determination.

**Figure 2.**
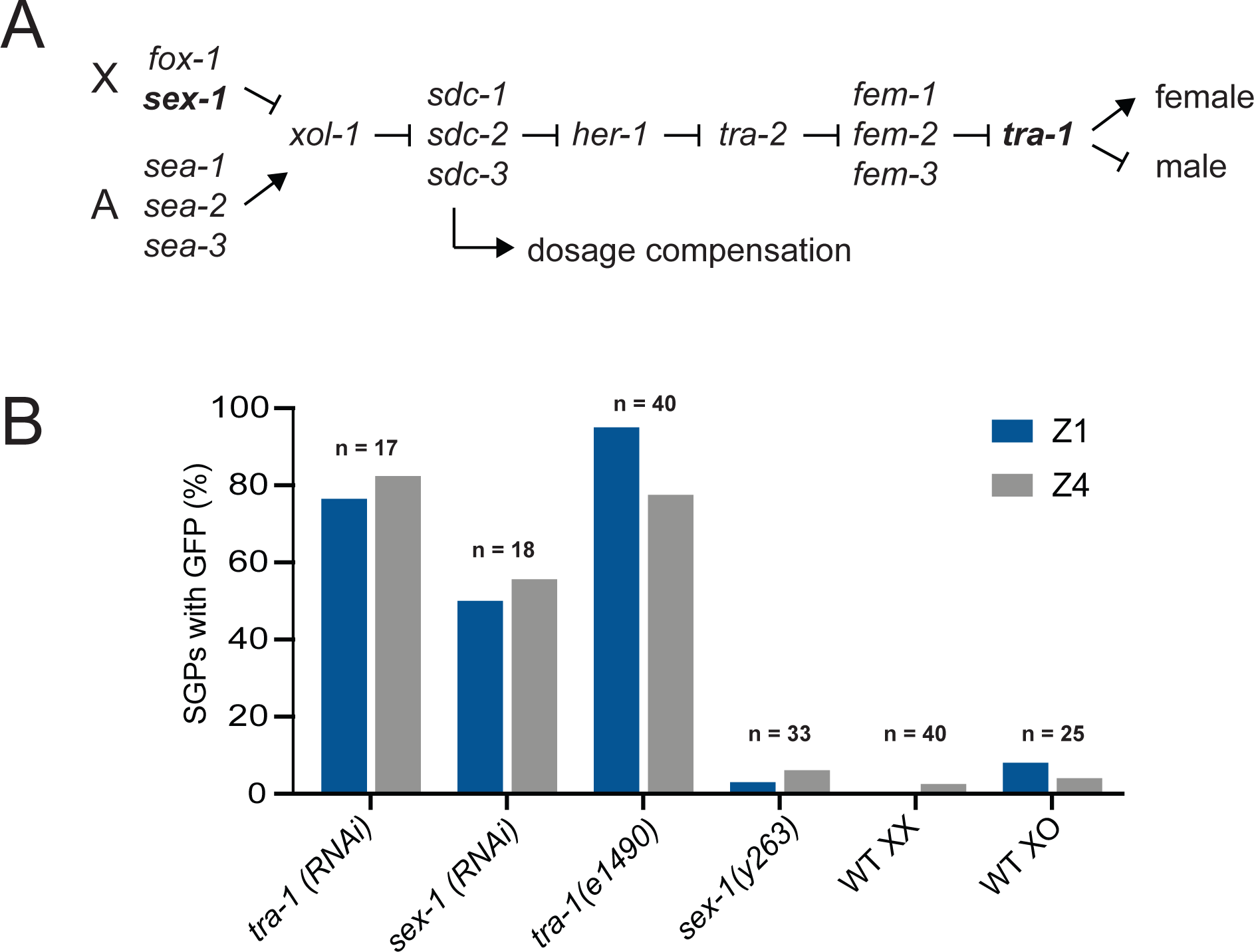
*tra-1* regulates the SGP/hmc cell fate decision. (A) The *C. elegans* sex determination pathway. Arrows indicate positive regulatory relationships; bars indicate negative regulatory relationships. The ratio of the number of X chromosomes to autosomes (X:A) determines the activity of *xol-1* which is transmitted through a series of negative regulatory interactions to ultimately regulate the activity of *tra-1*. The *sdc* genes also regulate dosage compensation. *sex-1* and *tra-1* (bold) lie at opposite ends of the pathway. (B) Expression of *arg-1::GFP* in SGPs of *tra-1* and *sex-1* RNAi or loss-of-function mutants (XX animals) compared to wild-type XX hermaphrodites and XO males.

We performed a secondary screen of all 28 candidate SGP fate regulators. The primary screen found differences in both the percentage of SGPs with *arg-1::GFP* expression and the level of *arg-1::GFP* expression in SGPs (Table 2). Therefore, in order to better classify our candidate SGP regulators, we recorded three levels of *arg-1::GFP* expression in SGPs – dim (as seen in the control), distinct, and bright (Figure 3). We included *pbrm-1*, one of four genes known to affect the SPG/hmc cell fate decision (Large and Mathies, 2014), as a positive control. *pbrm-1* is not an SGP-biased TF because it is expressed in both SGPs and hmcs (Mathies *et al*., 2019). In control worms, we observed a small percentage of SGPs with dim GFP expression (7.4%, *n* = 122 SGPs). By contrast, *pbrm-1(RNAi)* resulted in 45% of SGPs expressing GFP and most had distinct or bright expression. RNAi of the candidate SGP fate regulators resulted in a wide distribution of GFP expression in SGPs (Figure 3). Only two gene knockdowns, *tra-1* and *swsn-3,* had a higher percentage of SGPs with GFP expression than *pbrm-1;* both had distinct or bright expression. Interestingly, *swsn-3* encodes a subunit of SWI/SNF complexes that is predicted to be in a complex with PBRM-1. Of the remaining gene knockdowns, six had more than 25% of SGPs expressing *arg-1::GFP*, supporting the idea that the SGP/hmc cell fate decision is regulated by multiple TFs. In contrast to the multiple TFs which when depleted increased hmc marker expression, we saw only minor effects on SGP marker expression. Control animals always had two SGPs with *ehn3-tdTomato* expression, as did almost all of the RNAi knockdowns (File S1). The four exceptions were *mig-5, sex-1, nhr-136,* and *nhr-91,* which each had a single worm out of ∼40 with one or no *tdTomato*-expressing cells in the gonad; these worms had poor morphology by DIC making it difficult to assess the cause of the expression difference. In addition, we noted that *tra-1(RNAi)* resulted in worms with both SGPs positioned at the anterior end of the gonad; this phenotype is seen in *tra-1* loss-of-function mutants and results from the migration of Z4 toward the anterior of the gonad prior to its division (Mathies et al., 2004).

**Figure 3.**
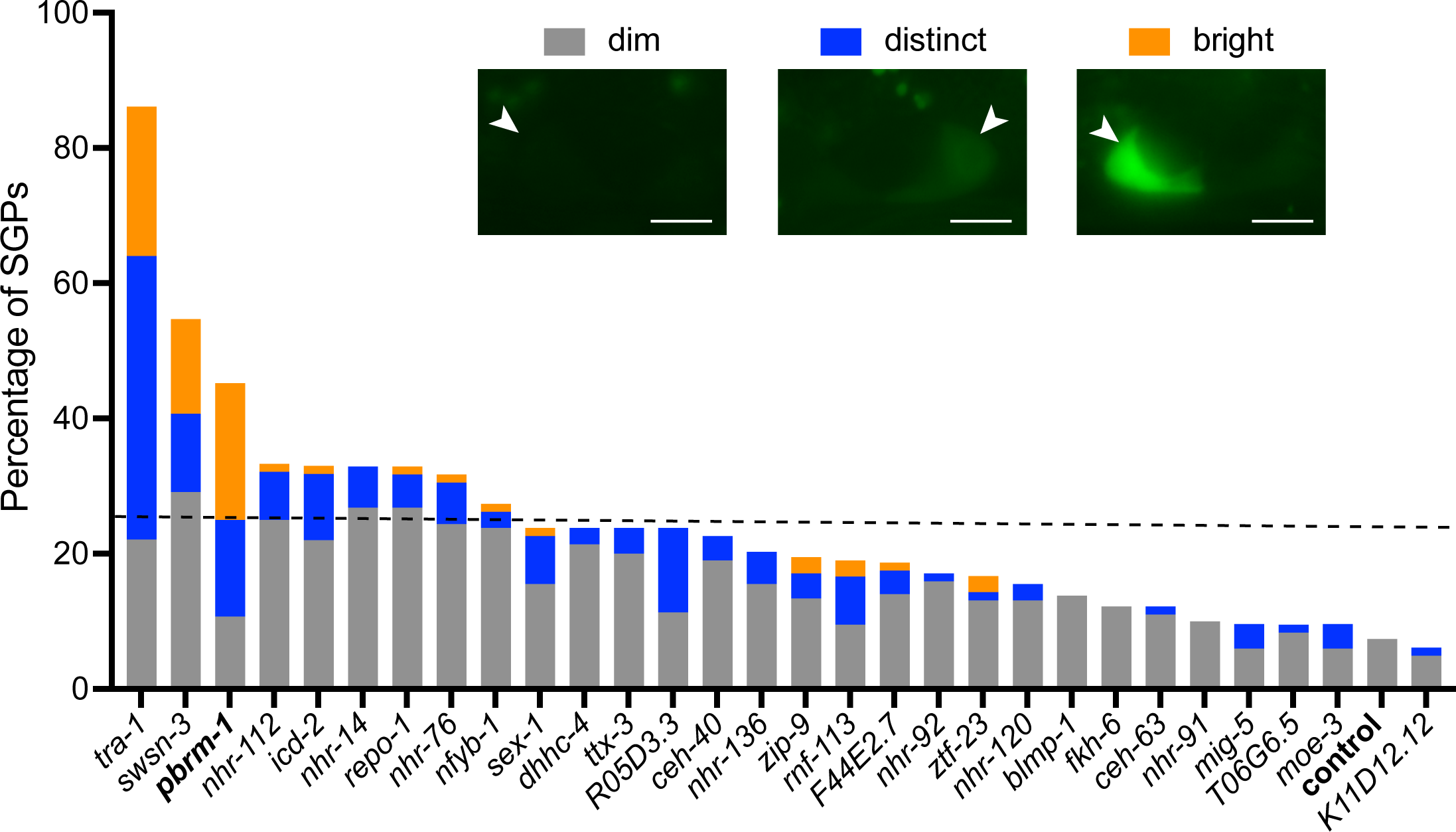
Genes affecting the expression of an hmc marker in SGPs Expression of *arg-1::GFP* in SGPs. The gene being inactivated by RNAi is indicated on the X-axis; *pbrm-*1 and the empty vector control are in bold. The percentage of SGPs expressing dim (gray), distinct (blue), or bright (orange) GFP in SGPs is on the Y-axis. The dotted line indicates our cutoff at ≥ 25% of SGPs with GFP expression. Insets show examples of dim, distinct, and bright expression. All images are 1.5 ms exposure with identical adjustment. Dim expression fades quickly and is difficult to capture. Scale bar is 5 μm.

Taken together, our results indicate that several SGP-biased TFs are important for the SGP cell fate. These data further suggest that determination and/or maintenance of the multipotent SGP cell fate is driven by repression of the hmc terminal cell fate.

### L4 RNAi screen identifies candidate multipotency factors

We screened L4 staged worms using a dissecting microscope and looked for abnormalities in gonadal morphology, including missing gonad arms, disorganized gonads, and absent gonads. Control RNAi worms always had a normal gonad (Table 3).

**Table 3.**
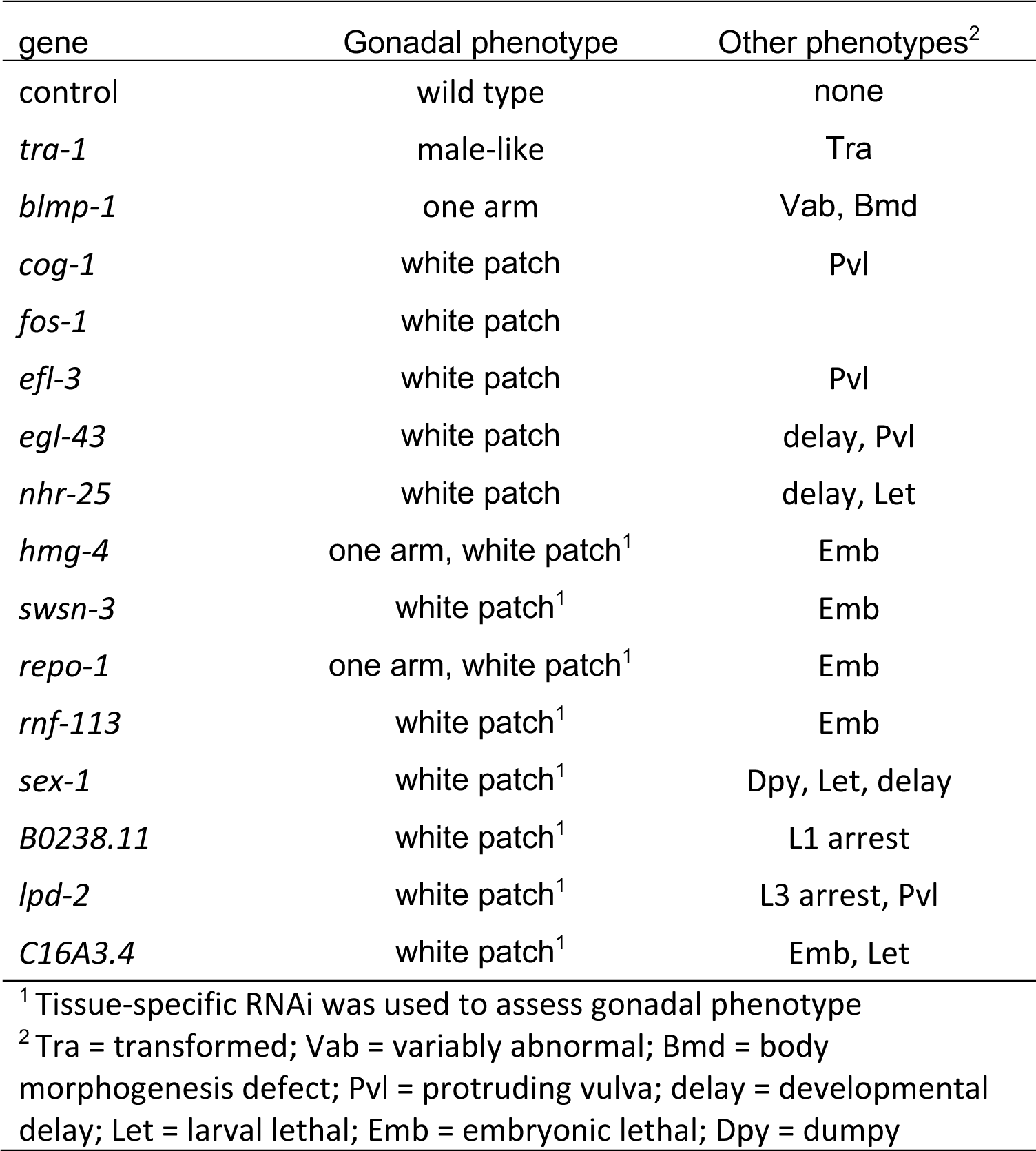
SGP-biased TFs with gonadal phenotypes.

Several RNAi knockdowns resulted in worms with a ventral “white patch” under the dissecting microscope. This phenotype is due to the presence of excess gonadal tissue near the vulva; it can result from a variety of somatic gonadal defects, including a failure to produce DTCs as is seen in Wnt pathway mutants (Siegfried et al., 2004; Siegfried and Kimble, 2002). RNAi knockdown of three genes, *blmp-1*, *hmg-4,* and *repo-1*, resulted in worms with missing gonadal arms. This phenotype can result from the absence of SGPs or the failure of SGPs to develop into gonadal arms (Large and Mathies, 2010; 2014; Mathies *et al*., 2003). RNAi knockdown of *tra-1* resulted in a male-like gonad, as expected for *tra-1* loss-of-function (Hodgkin, 1987). In total, our L4 screen identified 15 candidate SGP regulators (Table 3), six of which overlapped with our L1 screen.

In order to more specifically identify genes that regulate the proliferation and developmental potential of SGPs, we performed a secondary screen using a GFP reporter that is expressed in all descendants of the SGPs (Shaffer and Greenwald, 2022). We examined all genes that produced a phenotype in our L4 screen (Table 3), with the exception of *tra-1,* because the worms are transformed into males and the number of SGP descendants and organization of the gonad is different in males and hermaphrodites (Kimble and Hirsh, 1979). We screened five additional genes, *icd-2*, *nhr*-*14*, *nhr-76*, and *nfyb-1,* that had significant expression of the hmc marker in SGPs at the L1 stage (Figure 3) but appeared wild type in our L4 screen. We reasoned that these RNAi knockdowns might result in differences in the number or identity of SGP descendants that did not cause gross morphological defects at the L4 stage.

We examined the worms early in the L3 larval stage, when the hermaphrodite gonad contains 12 somatic cells, and we counted the number of GFP-expressing cells in the gonad (Figure 4A-B; File S1). Control worms always had 12 SGP descendants and the cells were correctly organized, with two DTCs at the poles of the gonad and the remaining 10 cells centrally localized in the SPh (Figure 4B-D). Nine RNAi knockdowns also had 12 SGP descendants that formed an SPh, suggesting that, although these genes produced a phenotype at the L1 or L4 stage, they are unlikely to be important for SGP proliferation or multipotency.

**Figure 4.**
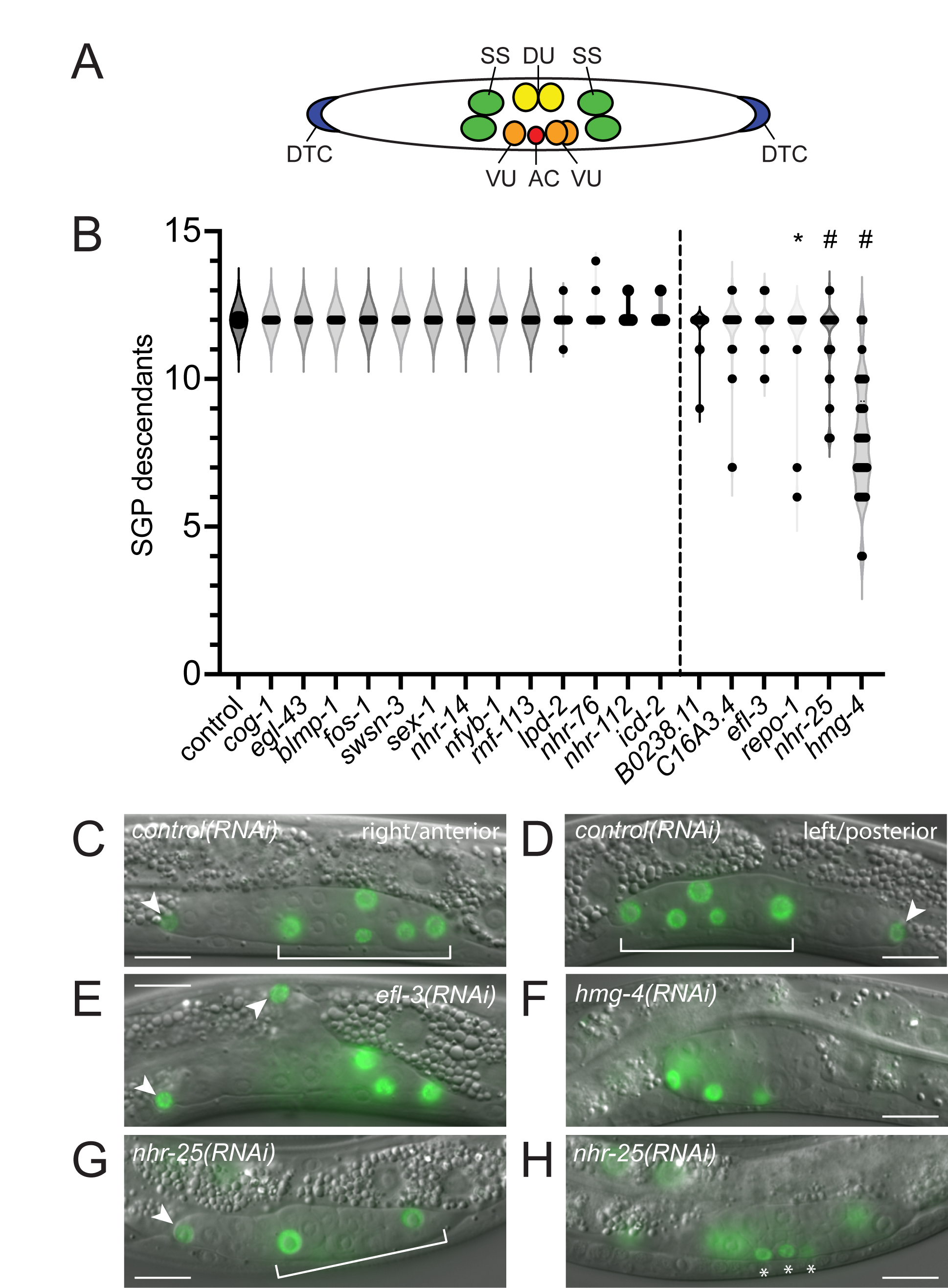
Genes affecting the number of SGP descendants at the L3 stage. (A) Organization of the somatic gonad in early L3 larvae. Anterior is to the left; ventral is down. Distal tip cells (DTCs, blue) are located at the anterior and posterior poles of the gonad. Remaining somatic gonadal cells are centrally located with sheath/spermathecal (SS, green) precursors closest to DTCs, dorsal uterine (DU, yellow) and ventral uterine (VU, orange) precursors more centrally located, and the anchor cell (AC) located at the center of the gonad. (B) Violin plot of the number of SGP descendants in early L3 larvae. The gene being inactivated by RNAi is indicated on the X-axis. Statistical comparisons were made using one-way ANOVA with Dunnett’s post-hoc tests to compare each RNAi to the control. Statistical significance is indicated; * p ≤ 0.05; # p ≤ 0.0001. (C-H) GFP fluorescence image overlay onto DIC images. GFP exposures are 200 ms. Scale bar is 10 μm. Arrowheads indicate DTCs; brackets indicate the somatic primordium (SPh) when present. (C-D) Empty vector RNAi control. The anterior arm (C) is on the right side of the worm; posterior arm (D) is on the left side of the worm. SPh is formed normally. (E) *efl-3(RNAi)* results in extra DTCs that appear to develop in place of the dorsal SS cell. (F) *hmg-4(RNAi)* results in fewer SGP descendants, no DTCs, and a round, underdeveloped gonad. Four SGP descendants are visible; one is out of the plane of focus. (G-H) *nhr-25(RNAi)* results in worms with fewer central SGP descendants (G) and several small cells that are not organized into the SPh (asterisks).

RNAi targeting three genes, *hmg-4, nhr-25,* and *repo-1*, caused a significant difference in the number of SGP descendants. The most extreme phenotype was seen in *hmg-4(RNAi)*, which had between four and twelve SGP descendants (8.0 +/- 1.9, *n* = 46). The SGP daughter cells were typically located at the periphery of the developing gonad (Figure 4F), and DTCs were frequently absent. These observations are consistent with the ventral white patch phenotype that we observed at the L4 stage. *nhr-25(RNAi)* worms had between 8 and 13 descendants, and they displayed two predominant phenotypes. First, they had two DTCs with fewer central cells (Figure 4G). Second, they had one or no DTCs, often accompanied by smaller cells that were not organized into the SPh (Figure 4G-H). These phenotypes are distinct from the previously described phenotype for *nhr-25(RNAi)* (Asahina *et al*., 2006; discussed below). *repo-1(RNAi)* had a low percentage of worms with six or seven GFP-positive cells in the gonad (5.9%, *n* = 51); these worms appeared to have descendants of only one SGP, judging by the positions and sizes of the daughter cells. We always observed two SGPs in *repo-1(RNAi)* L1s, suggesting that one of the two SGPs failed to proliferate and produce the appropriate cell types. This phenotype is reminiscent of *hnd-1* or SWI/SNF mutants (Large and Mathies, 2014). Finally, although it did not have a statistically significant difference in the number of SGP descendants, *efl-3(RNAi)* produced a highly penetrant and distinct phenotype. *efl-3(RNAi)* worms almost always had three or four DTCs in the gonad (89.1%, *n* = 46); the extra DTCs were often seen in worms with 12 SGP descendants and appeared to result from a transformation of SS precursors into DTCs (Figure 4E). Thus, our screen identified at least four genes that are important for generating the correct number and type of SGP descendants at the L3 stage, only one of which, *nhr-25*, had previously defined functions in the somatic gonad.

### *nhr-25* and *efl-3* are important for generating the correct number of DTCs

The first division of the SGPs establishes the proximal-distal axis of the gonad: central daughters adopt proximal fates and produce AC/VU precursors, while distal daughters adopt distal fates and produce DTCs (Figure 5A). Wnt signaling and *nhr-25* act in opposition to determine proximal and distal fates in the gonad (Asahina *et al*., 2006; Siegfried *et al*., 2004; Siegfried and Kimble, 2002), with the Wnt pathway promoting distal fates and *nhr-25* promoting proximal fates. Full proximal-to-distal cell fate transformation in *nhr-25(RNAi)* resulted in four DTCs and no AC (Asahina *et al*., 2006). In contrast to this published report, we did not observe extra DTCs in *nhr-25(RNAi)*; in fact, we often observed fewer than two DTCs. Interestingly, RNAi was performed differently in the two experiments. The previous study used feeding RNAi starting with newly hatched L1 larvae (larval RNAi), whereas we used feeding RNAi beginning with the parental generation and continuing through the L3 stage (systemic RNAi). Systemic RNAi should reduce *nhr-25* mRNA at an earlier time in development, which may account for the earlier and distinct phenotype we observed.

**Figure 5.**
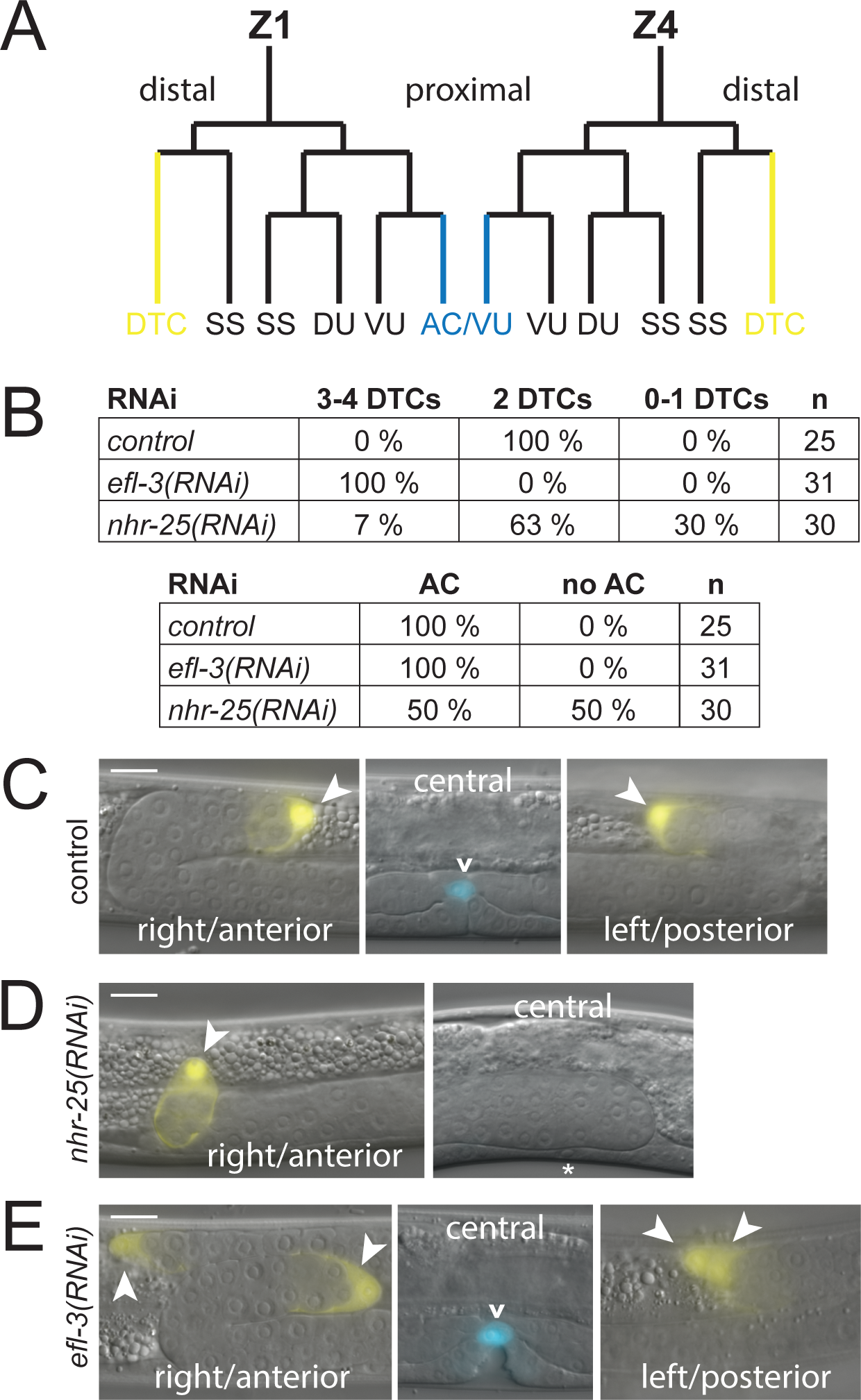
*nhr-25* and *efl-3* are required for the correct number of DTCs. (A) SGP cell (Z1 and Z4) lineages. Each SGP produces six cells by the end of the L2 larval stage. The first division of Z1 and Z4 produces distal and proximal daughter cells with different fates: proximal daughters (Z1.p and Z4.a) make dorsal and ventral uterine precursors (DU and VU), a sheath/spermathecal precursor (SS), and a cell that can become the anchor cell (AC/VU, blue); distal daughters (Z1.a and Z4.p) make a distal tip cell (DTC; yellow) and an SS cell. One of the AC/VU precursors becomes the AC; the cell that will adopt the AC fate is variable from animal to animal and determined by lateral signaling. (B) Number of DTCs and ACs in control, *nhr-25,* and *efl-3* RNAi; n = number of worms. (C-E) Expression of *lag-2::YFP* (yellow) and *cdh-3::CFP* (blue) fluorescence. Fluorescent image overlay on DIC image. Left panel (right side, anterior); middle panel (central plane); right panel (left plane, posterior). YFP exposures are 200 ms; CFP exposures are 300 ms. Scale bar is 10 μm. Arrowheads indicate DTCs; carat indicates the AC. (C) Control RNAi worms had two DTCs at the leading end of the anterior and posterior gonadal arms and one AC above the developing vulva. (D) *nhr-25(RNAi)* worms often had only one DTC at the leading end of a single gonadal arm (here, the anterior arm); they also frequently did not have an AC. Uninduced vulval precursor cell (asterisk). (E) *efl-3(RNAi)* worms always had additional DTCs. Here, four DTCs sit at the ends of gonadal arms; two DTCs anterior and two DTCs posterior. Extra DTCs were present in worms with an AC.

To examine the phenotype of systemic *nhr-25(RNAi)* in more detail, we used a strain that contained markers for DTCs (*lag-2::YFP*) and the AC (*cdh-3::CFP*). We examined the worms as early L4 larvae, when both markers are expressed in the appropriate cell type. Control worms always had two DTCs and one AC (Figure 5B-C). By contrast, *nhr-25(RNAi)* worms had between 0 and 4 DTCs. The most common phenotype was a single gonad arm led by one DTC (Figure 5D). Approximately half of the time, *nhr-25(RNAi)* worms had no AC as assessed by marker expression and the absence of vulval induction. The absence of the AC was independent of the number of DTCs, suggesting that it was not simply the result of a proximal-to-distal cell fate transformation. We conclude that *nhr-25* is required for SGPs to produce the correct number and array of cell types, in addition to its previously defined role in specifying proximal gonadal cell fates.

To confirm that *efl-3(RNAi)* results in ectopic DTCs, we examined *efl-3(RNAi)* using the AC and DTC markers. We found that *efl-3(RNAi)* always resulted in extra DTCs and a single AC (Figure 5B). Most of the worms had four DTCs and one AC (Figure 5E), indicating that the extra DTCs did not result from proximal-to-distal cell fate transformation in the first SGP division. Instead, the extra DTCs appeared to result from the transformation of SS cells into DTCs, as evidenced by their location on the dorsal surface of the gonad, which is the normal location of SS precursors (Figure 5E).

## Discussion

In this study, we conducted screens for regulators of cell fate and multipotency in a model multipotent lineage. We used RNAi to identify SGP-biased transcription factors that are important for the fate and/or developmental potential of SGPs. Seven genes were required for the expression of appropriate cell fate markers in SGPs but were dispensable for their subsequent development; these genes are likely to be involved in the determination or maintenance of the SGP fate. Three genes were required for the generation of the correct number and/or type of SGP descendants but did not have abnormal marker expression in SGPs; these genes are likely to include those that regulate the proliferation and/or developmental potential of SGPs, both of which are hallmarks of multipotency. Only one gene affected both aspects of development, indicating that the regulation of cell fate and developmental potential are genetically separable.

### Redundancy in the regulation of the SGP/hmc cell fate decision

We previously identified HND-1 and components of the SWI/SNF chromatin remodeling complex as regulators of the SGP/hmc cell fate decision. Strong loss-of-function alleles of *hnd-1* or *pbrm-1,* a component of SWI/SNF complexes, displayed two incompletely penetrant phenotypes: 1) SGPs express markers for both the SGP and hmc cell fate, and 2) one or both of the gonadal arms fails to develop (Large and Mathies, 2014; Mathies *et al*., 2003). We interpret these phenotypes as being the result of a partial cell fate transformation of SGPs into hmcs. Importantly, no single mutation, including strong loss-of-function and probable null alleles, is capable of fully transforming SGPs into hmcs, arguing for redundancy in the regulation of the SGP/hmc cell fate decision. Here we identified eight additional genes for which RNAi knockdown resulted in expression of both SGP and hmc cell fate markers in SGPs.

Only one gene, *repo-1,* was required for normal expression of cell fate markers in SGPs and for the development of two gonadal arms. Therefore, *repo-1* falls into the same phenotypic class as *hnd-1* and SWI/SNF genes. *repo-1* encodes a homolog of SF3a66 which is a component of U2 snRNPs and is involved in pre-mRNA splicing (Takenaka et al., 2004). Very little is known about the function of *repo-1* in *C. elegans.* A semi-dominant allele of *repo-1* causes reversed polarity in the early embryo and loss-of-function alleles result in embryonic lethality without a polarity defect (Keikhaee et al., 2014). More recently, *repo-1* was found to be important for longevity and, in this context, it is thought to function as a splicing factor (Heintz et al., 2017). *repo-1* was included on the TF2.0 transcription factor list because of its C2H2 zinc finger domain. C2H2 zinc fingers can bind RNA or DNA (Cassandri et al., 2017); therefore, it remains to be seen if REPO-1 acts as a transcription factor or a splicing factor during SGP development.

Interestingly, most of the RNAi knockdowns that altered expression of cell fate markers in SGPs (Figure 3) had little or no effect on the number of SGP descendants (Figure 4). This observation strongly suggests that SGPs can recover from partial cell fate transformation to produce a normal somatic gonad. This is exemplified by *swsn-3(RNAi),* which produced strong ectopic expression of the hmc marker in SGPs yet had no effect on the number or type of SGP descendants at the L3 stage. *swsn-3* encodes an accessory subunit of the SWI/SNF chromatin remodeling complex; therefore, it is predicted to function with *pbrm-1, swsn-1,* and *swsn-4* (Large & Mathies, 2014). The biochemistry of the SWI/SNF complex has not been worked out in *C. elegans.* One possibility is that SWSN-3 is only present in SWI/SNF complexes in SGPs and not later during cell lineage progression. Alternatively, the effectiveness of RNAi may have been influenced by background differences between the strains used to assess these two phenotypes. The only existing *swsn-3* allele is viable and has no effect on gonadogenesis (Large and Mathies, 2014), but because *swsn-3(RNAi)* causes embryonic lethality, it seems likely that this allele does not cause a strong loss of function. Clarification of the roles of *swsn-3* in somatic gonadogenesis will have to await the isolation of a strong *swsn-3* allele.

### *tra-1/GLI* is important for distinguishing SGPs from hmcs

A majority of *tra-1* mutant worms strongly expressed the hmc marker in SGPs, indicating that *tra-1* is required to suppress gene expression characteristic of a terminally differentiated cell in the multipotent SGPs. This expression difference cannot be explained by the sexual transformation of XX animals into males in *tra-1* mutants because wild type (XO) males do not express *arg-1::GFP* in SGPs. We previously showed that *tra-1* regulates symmetry in the gonad primordium, independent of its role in sex determination (Mathies *et al*., 2004), and that this function is conserved in other nematode species (Kelleher et al., 2008). We have argued that these non- sex-specific functions of *tra-1* might represent the more ancestral function of *Ci*/*Gli* genes. First identified in *Drosophila*, Ci acts as the terminal transcription factor in the Hedgehog (Hh) signal transduction pathway and is important for patterning in the embryo and imaginal discs (reviewed in Huangfu and Anderson, 2006). Three vertebrate *Gli* genes have overlapping functions downstream of Sonic hedgehog (Shh) signaling and are important for central nervous system patterning and lung development (reviewed in Matise and Joyner, 1999). In this light, the involvement of *tra-1* in the SGP/hmc cell fate decision might represent another example of *tra-1*’s more ancestral functions.

### HMG-4 and the maintenance of cell fate

*hmg-4* encodes a subunit of the *C. elegans* FACT (facilitates chromatin transcription) complex. FACT is composed of two subunits, SSRP1 (structure-specific recognition protein 1) and SPT16 (Suppressor of Ty 16), and was identified for its role in promoting transcript elongation through nucleosomes (Orphanides et al., 1998; Orphanides et al., 1999). *C. elegans* FACT is required for normal cell cycle timing during embryogenesis (Suggs et al., 2018) and acts as a barrier to cellular reprogramming in adult tissues (Kolundzic et al., 2018). We found that *hmg-4(RNAi)* resulted in a reduced number of SGP descendants at the L3 stage and abnormal gonadal morphology at the L4 stage. Based on the absence of gonadal arm elongation, we infer that at least one cell type, the DTC, was absent in *hmg-4(RNAi)* worms. Therefore, *hmg-4* function is necessary for SGPs to generate the correct number and type of descendants, suggesting a role in SGP proliferation and multipotency. FACT functions as a barrier to reprogramming, in part, by maintaining cell fate. One possibility is that FACT maintains the SGP cell fate and, in its absence, the cells fail to execute their developmental program. FACT is also required for transcript elongation and results in longer cell cycle times. Alternatively, *hmg-4* might be required for proper execution of the cell division cycle during SGP development resulting in fewer SGP daughter cells.

### EFL-3 and cell fate

*efl-3* is one of three E2F-enconding genes in the *C. elegans* genome. It is most similar to two mammalian genes, E2F7 and E2F8, and was identified for its ability to inhibit cell death in ventral cord neurons by promoting expression of the pro-apoptotic gene *egl-1* (Winn et al., 2011). It is also expressed in seam cell lineages and is required for development of the correct number of seam cells in the adult (Katsanos et al., 2021). We show here that loss of *efl-3* function in the somatic gonad results in one or two additional DTCs. The supernumerary DTCs appeared to develop at the expense of the dorsal SS precursor, based on their position within the developing gonad and on the absence of dorsal SS cells in gonads with extra DTCs. The dorsal SS precursor is the sister of the DTC; therefore, one simple explanation is that *efl-3* is required for specification of the SS cell fate and/or inhibition of the DTC fate. E2F proteins are important regulators of the cell cycle, acting with DP to promote cell proliferation (Lam and La Thangue, 1994).

EFL-3 and its homologs, E2F7 and E2F8, lack important transactivation and interaction domains typically found in more canonical E2F proteins (Lammens et al., 2009).

However, there is evidence to suggest that atypical E2F proteins act in cell cycle regulation. For example, human E2F7 promotes proliferation and inhibits differentiation of acute myeloid leukemia (AML) cells (Salvatori et al., 2012) and E2F8 overexpression promotes proliferation and tumorigenicity in breast cancer cell lines (Salvatori *et al*., 2012). SS cells continue to divide and generate multiple cell types, whereas DTCs are terminally differentiated. Therefore, another possibility is that EFL-3 is acting to promote proliferation in SS cells and that, in its absence, the cells differentiate into DTCs.

### *nhr-25* is a pleiotropic regulator of somatic gonad development

The first division of the SGPs is asymmetric and is governed by a Wnt/μ-catenin signaling pathway that promotes distal fates. Mutations in Wnt pathway mutants result in all SGP daughter cells adopting a proximal fate; no DTCs are produced and the gonad does not elongate (Siegfried *et al*., 2004; Siegfried and Kimble, 2002). *nhr-25* opposes Wnt signaling such that *nhr-25* loss-of-function results in extra DTCs at the expense of proximal cells (Asahina 2006). We found that *nhr-25* RNAi, when applied at an earlier stage in development, resulted in fewer DTCs and a loss of the AC. The simplest explanation for these different observations is that *nhr-25* acts early in SGPs and then again in opposition to Wnt signaling to determine proximal and distal fates. *nhr-25* acts in the epidermis to promote cell differentiation by repressing the expression of factors that promote a stem cell fate (Katsanos and Barkoulas, 2022). It is hard to reconcile the *nhr-25* phenotype in the somatic gonad with a role in promoting differentiation. Instead, our results suggest that *nhr-25* promotes multipotency in SGPs: *nhr-25* loss of function does not affect the SGP cell fate but does affect the number and type of cells produced by SGPs. It will be interesting to compare the function of *nhr-25* in multipotent SGPs with that in multipotent epidermal stem cells.

### Data Availability

Strains are available upon request. File S1 contains all primary data from the RNAi screens.

## Acknowledgements

The authors thank Justin Shaffer and Iva Greenwald for sharing strains prior to publication and David Bilder for critical reading of the manuscript. Strains were provided by the *Caenorhabditis* Genetics Center and the *C. elegans* Reverse Genetics Core Facility at the University of British Columbia, which is part of the international *C. elegans* Gene Knockout Consortium.

